# GeNNius: An ultrafast drug-target interaction inference method based on graph neural networks

**DOI:** 10.1101/2023.06.20.545544

**Authors:** Uxía Veleiro, Jesús de la Fuente, Guillemo Serrano, Marija Pizurica, Mikel Casals, Antonio Pineda-Lucena, Silve Vicent, Idoia Ochoa, Olivier Gevaert, Mikel Hernáez

**Affiliations:** Center for Applied Medical Research University of Navarra, IdiSNA, Pamplona, Spain; Department of Electrical and Electronics Engineering, School of Engineering (Tecnun), University of Navarra, Spain; Center for Data Science, New York University, New York; Stanford Center for Biomedical Informatics, Stanford University, California; Internet technology and Data science Lab (IDLab), Ghent University, Technologiepark-Zwijnaarde 126, Ghent, 9052, Gent, Belgium; Instituto de Ciencia de los Datos e Inteligencia Artificial (DATAI), University of Navarra, Pamplona, 31008, Spain

**Keywords:** drug repurposing, graph neural networks, drug-target interaction

## Abstract

Drug-target interaction (DTI) prediction is a relevant but challenging task in the drug repurposing field. In-silico approaches have drawn particular attention as they can reduce associated costs and time commitment of traditional methodologies. Yet, current state-of-the-art methods present several limitations: existing DTI prediction approaches are computationally expensive, thereby hindering the ability to use large networks and exploit available datasets and, the generalization to unseen datasets of DTI prediction methods remains unexplored, which could potentially improve the development processes of DTI inferring approaches in terms of accuracy and robustness. In this work, we introduce Gennius (Graph Embedding Neural Network Interaction Uncovering System), a Graph Neural Network (GNN)-based method that outperforms state-of-the-art models in terms of both accuracy and time efficiency across a variety of datasets. We also demonstrated its prediction power to uncover new interactions by evaluating not previously known DTIs for each dataset. We further assessed the generalization capability of Gennius by training and testing it on different datasets, showing that this framework can potentially improve the DTI prediction task by training on large datasets and testing on smaller ones. Finally, we investigated qualitatively the embeddings generated by Gennius, revealing that the GNN encoder maintains biological information after the graph convolutions while diffusing this information through nodes, eventually distinguishing protein families in the node embedding space.

**Code Availability:** https://github.com/ubioinformat/GeNNius

## 1. Introduction

The process of identifying new drugs to treat a specific disease can be simplified by seeking a chemical compound that modulates a pharmacological target implicated in that disease, with the goal of altering its biological activity. Even though different biological entities can be chosen as targets, such as RNA or proteins, the latter are the most common pharmacological targets (1). Targeting proteins allows the modulation of many biological processes implicated in maintaining health and potentially preventing or treating diseases. For example, drugs targeting metabolic enzymes can alter how cells process nutrients and energy (2).

Although high-throughput wet-lab techniques were developed to accelerate drug discovery pipelines, both in vitro and in vivo approaches are time-consuming and can be costly (3). To address these limitations, computational methods have arisen as promising tools to reduce the time and resources required to bring new treatments to market. The field of drug repurposing involves predicting novel drug-target interactions (DTIs) that will ultimately enable the discovery of new uses for already approved drugs (4). In recent years, the availability of large amounts of data has made it possible to design machine learning models that can assist in these drug development tasks, through, for example, the identification of complex molecular patterns that were not previously uncovered. These models typically leverage multiple types of data, including amino acid sequences (5) and the 3-D protein structures (6), as recent advances in protein structure prediction such as AlphaFold (7, 8) have significantly increased the amount of structural information available.

Specific to DTI prediction, several different machine learning architectures have been proposed in recent years (see Methods). While some models utilize simple linear regression techniques, others contain more complex mechanisms such as transformers (9). However, most of these technologies do not consider the global view of how proteins and drugs are connected, which could be informative towards the discovery of novel relationships. To allow for modeling the network topology, recent works have been proposed to represent DTI data as a graph (10, 11). Specifically, DTIs can be modeled as a heterogeneous graph connecting drugs and proteins (both represented as nodes) based on recorded interactions in wet-lab experiments (edges). This representation can be augmented by adding additional node types (e.g., diseases), or edge types (e.g., protein similarity). The DTI prediction model is then trained to predict whether a drug has the potential to interact with a protein.

Advances in machine learning for graphs have highlighted Graph Neural Networks (GNNs) as a powerful tool to model these complex networks for a wide range of applications across diverse fields including economics (12), particle physics (13), and especially biomedicine (14). The defining characteristic of a GNN is that it uses a form of neural message passing, where at each iteration the hidden embeddings of the nodes are updated (15). Further, contrary to most previous node embedding techniques such as Node2Vec (16), GNNs are able to generalize from a set of training examples to unseen data points. This capability is of utmost importance to guarantee the generalization capabilities of the developed models when facing unseen interactions.

Recently, entire libraries have been developed to work with GNNs. Special mention should be made to PyTorch Geometric (PyG), a geometric deep learning library built on top of PyTorch (17). Among other functions and layers, PyG implements the SAGEConv layer, which corresponds to the GraphSAGE operator that was originally designed to allow the training of GNNs in large networks (18). SAGEConv simultaneously learns the topological structure of the neighborhood of each node, as well as the distribution of the features of the nodes in the neighborhood.

In this work, we present a novel DTI prediction method, termed Gennius (Graph Embedding Neural Network Interaction Uncovering System), built upon SAGEConv layers followed by a neural network (NN)-based classifier. Gennius outperforms state-of-the-art DTI prediction methods across several datasets, not only in AUROC and AUPRC, but also in execution time. Since ensuring the capabilities of in silico drug repurposing approaches to find new interactions is of utmost importance, we also evaluated the ability of Gennius to predict true interactions reported as negative in a given dataset, yielding promising results. We further assessed the generalization capability of our model by training in one dataset and testing in a different one. This procedure mimics more realistically how the model would behave in a real-world setting.

Finally, while drug repurposing approaches based on complex machine learning models have eased the discovery of new targets, they often lack explainability. We analyzed qualitatively how drug features (such as commonly-used molecular descriptors) and protein features (such as the amino acid ratio of protein sequences) are combined in a non-linear manner while solving the DTI prediction task. This analysis revealed that the GNN encoder maintains biological information while diffusing this information through nodes, eventually distinguishing protein families in the node embeddings. Overall, the results of our evaluation provide strong support for the effectiveness of Gennius, and introduce relevant guidelines to build GNN-based drug repurposing approaches.

## 2. Materials and Methods

### 2.1. Methods

#### 2.1.1. Model architecture

GeNNius architecture is composed of a Graph Neural Network (GNN) encoder that generates node embeddings and a Neural Network (NN)-based classifier that aims to learn the existence of an edge (i.e., an interaction) given the concatenation of a drug and protein node embeddings (Figure 1).

**Fig. 1.**
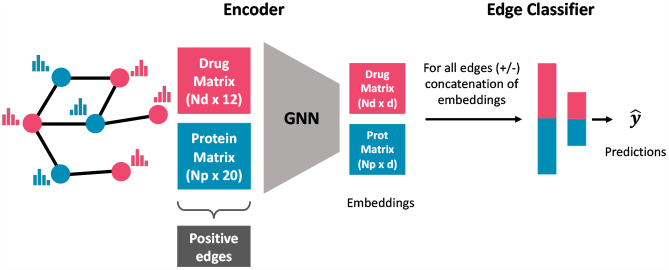
Schematic of Gennius architecture. Gennius inputs a graph containing drug (red) and protein (blue) nodes, where *N*_*d*_ and *N*_*p*_ represent the number of drugs and proteins, respectively. First, a GNN generates node representations with an embedding of dimension *d* = 17. Second, a NN-based classifier aims at learning the existence of an edge given a set of concatenations of drug and protein embeddings. Note that at this stage a negative set of edges is generated.

In GNNs, nodes in the graph exchange messages with their neighbors to update their feature representation, which is formulated with two fundamental functions: the message and the update functions.(19):

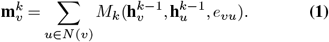

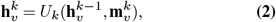

where *k* ∈ {1, …,*K*} represents the layer, **m**_*v*_ the aggregated message vector for node *v, N* (*v*) the neighbor nodes of *v*, and **h**_*v*_ ∈ ℛ^*d*^ the node *v* embedding, of dimension *d. M* (**h**_*v*_, **h**_*u*_, *e*_*vu*_) defines the message between node *v* and its neighbor node *u*, which depends on the edge information *e*_*vu*_. Finally, *U*_*k*_ is the node update function, which combines aggregated messages from the node’s neighbors with the node’s own representation.

Gennius’s encoder is composed of four SAGEConv layers, which are responsible for generating network-preserving node embeddings **h** ℛ^*d*^ (*d* = 17 in our case) by aggregating information from the embeddings of each node’s local neighborhood. Thus, in Gennius, the embedding of node *v* at SAGEConv layer *k* is given by:

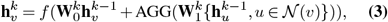

where *f* is the activation function (Tanh in our case) and AGG represents the aggregation function (SUM in our case). 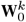 and 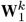 are the learnable weight matrices; since we are working with heterogeneous graphs, where a drug is only connected to proteins and vice versa, if 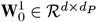 then 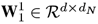, or the other way around, being *d*_*P*_ (*d*_*N*_) the initial dimension of proteins (drugs) node features. For *k>* 1, both matrices have dimension *d* ×*d*.

The NN-based classifier is composed of two dense layers, both using ReLu as the activation function, followed by the output layer, which is composed of a single neuron with a sigmoid activation function. The input to the classifier is a vector of dimension 2*d* (corresponding to the concatenation of a drug and protein embeddings), and the output is the estimated probability of having an interaction (positive edge). Gennius architecture (depicted in Figure 1) and hyperparameters were chosen through a grid search with ten independent runs, using different types and number (*K*) of GNN layers, different embedding dimension *d*, activation functions, aggregation functions, and different number of heads for layers with attention. This approach helped us to fine-tune the model (see Supplementary Material 1 for a detailed description of the process and hyperparameters).

#### 2.1.2. Model configuration

The model was trained with the Adam optimizer (20) and a learning rate of 0.01. We use a loss that combines the sigmoid of the output layer and the binary cross entropy in a single function. This combination takes advantage of the log-sumexp trick for numerical stability (21). Given a dataset divided into batches of size *N*, the loss *l*_*n*_ for sample *n* in the batch is computed as follows:

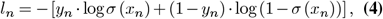

where *y*_*n*_ ∈ {0, 1 } is the associated label for sample *n, ŷ*_*n*_ = *σ*(*x*_*n*_) the estimated probability of the sample belonging to the positive class (i.e., existence of an interaction), and *x*_*n*_ the output of the last linear layer (before the activation function). The final batch loss *L* is then computed as the average of (*l*_1_,…, *l*_*N*_). Finally, a dropout of 0.2 is used at the encoder stage (the GNN) to address potential collinearities of node features (22) (dropout rate chosen through hyperparameter-tuning, see Supplementary Material Section 1).

The model is implemented with early stopping, calculated on validation data, with a minimum of 40 training epochs. The latter is especially useful for small datasets where an early stop may occur during the first epochs, eventually causing underfitting. The model was built with the latest version of PyTorch Geometric (2.3.0), with PyTorch 2.0.0-cuda11.7, and the following packages: pyg-lib (0.2.0+pt20cu117), torch-scatter (2.1.1+pt20cu117) and torch-sparse (0.6.17+pt20cu117). A Dockerfile for running the model is available at https://github.com/ubioinformat/GeNNius.

#### 2.1.3. Model training and evaluation

In the standard setting in which a single dataset (graph in our case) is used to evaluate model performance, the input graph is randomly split into a 70:10:20 ratio for train, validation, and test, respectively, via the random link split function of PyG. This function also randomly selects the negative edges needed for training and testing the NN-based classifier for a 1:1 positive/negative ratio. The training set requires further shuffling of positive and negative edges. Only 70% of train edges is used for training the encoder, while the rest are kept apart for the edge prediction part (i.e., the edge classifier).

To assess the performance of the models in the edge classification task on test data, we use the area under the Receiver Operating Characteristic curve (AUROC), as well as the area under the precision-recall curve (AUPRC), both widely used for evaluating DTI prediction models. We refer to Supplementary Material Section 2 for a more extended description of these metrics.

#### 2.1.4. Node features

Due to the different nature of drugs and proteins, we choose a vastly different set and dimension of features for drug and protein nodes. The protein node features are encoded as a 20-dimensional vector, accounting for the 20 different amino acids, where each feature indicates the proportion of the corresponding amino acid in the protein sequence associated to the node. Drug node features are chosen to be wellknown molecular descriptors, calculated with RDKit (23), from their SMILES. Specifically, the 12 selected features for drug nodes are: LogP value, molecular weight, number of hydrogen bond acceptors, number of hydrogen bond donors, number of heteroatoms (i.e., any atom that is not carbon or hydrogen), number of rotatable bonds, topological polar surface area (surface sum over all polar atoms, primarily O and N, also including their attached H atoms), number of rings and aromatic rings, number of NHs and OHs, number of nitrogen and oxygen atoms, number of heavy atoms a molecule (atomic number > 1), and number of valence electrons. While some of the above-mentioned features are related, model learning and performance is not expected to deteriorate as a dropout layer was introduced to reduce the potential effect of features’ collinearity (the correlation matrices of drug/protein features are provided in Supplementary Material Section 3). While other node features could be considered, such as protein pre-computed embeddings, training the model with those features showed almost no increase in performance (Supplementary Material Section 4). In addition, the pre-computed protein embeddings were not available for all proteins, which led to a decrease in nodes, hindering the training process and making it impossible for the model to generalize when trained in small networks (Supplementary Material Section 4).

#### 2.1.5. Related work

In order to benchmark our proposed method Gennius, we focus on the latest DTI prediction models that have been shown to outperform previously developed models in their respective publications.

- **DTINet** (10). It considers a heterogeneous graph with four node types (drugs, proteins, side effects and diseases) and six edge types (DTIs, protein-protein interaction, drug-drug interaction, drug-disease association, protein-disease association, drug-side-effect association, plus similarity edges between drugs and proteins). After compact feature learning on each network drugs/proteins, it calculates the best projection of one space onto another using a matrix completion method, and then infers interactions according to the proximity criterion. We note that the large quantity of data required to run the method hampered its reproducibility, as the code for generating all these matrices was not available.
- **EEG-DTI** (11). EEG-DTI, which stands for endto-end heterogeneous graph representation learningbased method, also considers a heterogeneous network, where nodes and edges are the same as in DTINet (see above). The model first generates a lowdimensional embedding for drugs and proteins with three Graph Convolutional Networks (GCN) layers, and then concatenates these layers for drugs and proteins separately. Finally, it calculates the inner product to get the protein-drug score.
- **HyperAttentionDTI** (5). This method only requires the SMILES string for drugs and the amino acid sequence for proteins. Then, it embeds each character of the different sequences into vectors. The model is based on the attention mechanism and Convolutional Neural Networks (CNNs), in order to make DTI predictions.
- **Moltrans** (9). Its name stands for Molecular Interaction Transformer for predicting DTIs. As HyperAttentionDTI above, it needs the SMILES for drugs and amino acid sequences for proteins. Then, it makes use of unlabeled data to decompose both drug and nodes into high-quality substructures, to later create an augmented embedding using transformers. Due to this architecture, the model is able to identify which substructures are contributing more to the overall interaction between a protein and a drug.

### 2.2. Materials

#### 2.2.1. Datasets

In this work we selected various datasets that have been widely used for DTI prediction tasks:

- **DrugBank** (24). Drug-Target interactions collected from DrugBank Database Release 5.1.9. Its first release was in 2006, although it has had significant upgrades during the following years.
- **BioSNAP** (25). Dataset created by Stanford Biomedical Network Dataset Collection. It contains proteins targeted by drugs on the U.S. market from DrugBank release 5.0.0 using MINER (26).
- **BindingDB** (27). Database that consists of measured binding affinities, focusing on protein interactions with small molecules. The binarization of the dataset was done by considering interactions positive if their *K*_*d*_ was lower than 30. Data downloaded from Therapeutics Data Commons (TDC) (28).
- **Davis** (29). Dataset of kinase inhibitors with kinases covering >80% of the human catalytic protein kinome. The binarization of the dataset has been done considering as positive interactions with a *K*_*d*_ lower than 30. Data downloaded from Therapeutics Data Commons (TDC) (28).
- **Yamanishi et al**. (30). It is composed of four subsets of different protein families: Enzymes (E), IonChannels (IC), G-protein-coupled receptors (GPCR) and nuclear receptors (NR). Yamanishi dataset has been considered the golden standard dataset for DTI prediction and has been used in several published models (11, 31, 32). DTIs in this dataset come from KEGG BRITE (33), BRENDA (34), SuperTarget (35) and DrugBank.

For all considered datasets, we dropped those drugs and proteins from which SMILES or amino acid sequences could not be generated, yielding slightly smaller networks (see Supplementary Material Section 5).

Note that the above-mentioned datasets, with the exception of BindingDB and Davis, contain only positive samples, i.e., positive links in the network. Nevertheless, when choosing negative samples, we performed random subsampling to have a balanced dataset prior to training the model.

Datasets statistics are summarized in Table 1. These datasets were released in different years, and thus some drug-target interactions can be shared across datasets (See Supplementary Material Section 6).

**Table 1.** Dataset Statistics.

|  | DrugBank | BioSNAP | BindingDB | Davis | E | Yamanishi<br>GPCR | IC | NR |
| --- | --- | --- | --- | --- | --- | --- | --- | --- |
| Number of drugs | 6823 | 4499 | 3084 | 59 | 444 | 222 | 210 | 53 |
| Number of proteins | 4652 | 2113 | 718 | 218 | 660 | 94 | 203 | 25 |
| Total number of nodes | 11475 | 6612 | 3802 | 277 | 1104 | 316 | 413 | 78 |
| Total number of edges | 23708 | 13838 | 5937 | 673 | 2920 | 634 | 1471 | 86 |
| Sparsity (%) | 0.07 | 0.15 | 0.27 | 5.52 | 1.01 | 3.13 | 3.57 | 6.94 |
| # Connected components | 412 | 174 | 231 | 1 | 44 | 18 | 3 | 10 |
| Avg degree drug nodes | 3.47 | 3.08 | 1.93 | 11.41 | 6.58 | 2.86 | 7.00 | 1.62 |
| Avg degree protein nodes | 5.10 | 6.55 | 8.27 | 3.09 | 4.42 | 6.74 | 7.25 | 3.44 |

#### 2.2.2. Dataset configuration for inferring unknown positives

DTI datasets contain information from diverse sources, have been released in different years, and may be curated in various ways. As a result, negatively labeled edges in one dataset may be reported as positive in other datasets. We evaluate these unknown positive edges for each dataset to asses if Gennius can predict them (see Supplementary Section 6 for details on the number of these edges for each dataset). Importantly, we ensured that testing edges do not appear as negatives during training to asses how well Gennius predicts these specific interactions; we repeated the process ten independent times, enabling us to investigate the variability of the prediction depending on training edges, which is often not reported in DTI prediction models.

#### 2.2.3. Data leakage prevention during evaluation on unseen datasets

Contrary to previously proposed models, we assess the generalization capability of Gennius by training it on one dataset and testing it on another. For a fair assessment, it is necessary to ensure that there is no data leakage of DTIs between training and testing.

Let us consider two nodes that are present both in the training and test datasets. There are four possible scenarios for an edge connecting these nodes. A positive edge in both datasets is a clear example of data leakage from the train to the test set, as we already informed the model about that positive edge during training. Hence those repeated DTIs are removed during training. On the other hand, edges that appear in one dataset but not on the other one are kept. Keeping the negative edges in the training data makes sense from a usability perspective since a non-reported DTI in a given dataset does not necessarily mean that that pair does not interact, and we would like to test the capabilities of the model under this general scenario. Further, a negative edge may be shared in both datasets; however, since negative edges are randomly selected when generating the training and testing sets, the probability of picking the same edge in both datasets is very low. As an illustrative example, when using DrugBank for training and NR for testing, the probability of selecting the same negative edge is approximately 3*e*^−6^.

We performed five independent training runs on each dataset, i.e., randomly selecting each time a different set of edges for training the model. Next, for each trained model, we performed five independent testing runs. We report the average and standard deviation of the AUROC and AUPRC metrics, of the test set, across the total 25 runs per training-testing dataset pair.

#### 2.2.4. Protein and Drug Annotation

Protein family and enzyme annotation was retrieved from the ChEMBL database (release 31), as its family hierarchy is manually curated and according to commonly used nomenclature (36). Drug chemical annotation was generated using ClassyFire, an automated structural classifier of chemical entities (37). Annotation was used for coloring t-SNEs.

#### 2.2.5. Hardware

All simulations were performed on a server with 64 intel xeon gold 6130 2.1Ghz cores with 754Gb of RAM and a NVIDIA GeForce RTX 3080, driver version 515.43.04, with cuda 11.7. version

## 3. Results

### 3.1. Gennius outperforms state-of-the-art methods

The proposed model was run on the eight selected datasets with five independent runs. The resulting AUROC and AUPRC metrics on the test sets across all datasets, as well as running times (corresponding to train, validation and test), are presented in Figure 2 (see also Supplementary Material Section 7). Gennius returned AUROC and AUPCR performance close to 1 (>0.9) for large datasets, and while smaller datasets reported worse results, they are still compelling (>0.8 in almost all runs). NR, being the smallest one, achieved the worst results (>0.7). Additionally, the large datasets showed stable results, with a low standard deviation, across the five independent runs. Further, the model execution time was ultrafast for all datasets (less than a minute for the largest dataset). Note that the time variance in the large datasets is due to early stopping.

Next, we compared the performance of Gennius with previously proposed methods. Table 2 shows the performance results of Gennius and the state-of-the-art methods for both DrugBank and BIOSNAP, the largest standard DTI datasets. We focus on these datasets as they better characterize the current size of testable available drugs. Gennius outperformed all benchmarked methods in terms of AUROC and AUPRC. Importantly, the execution time is significantly reduced, even when executed without GPU (see Supplementary Material Section 8). Previous methods’ running time was in the order of tens of minutes (except DTINet, which took 4.23 min), while Gennius took less than 0.6 minutes to perform the training, validation, and testing. The closest performance in AUROC and AUPRC to Gennius was achieved by EEG-DTI. However, EGG-DTI took four orders of magnitude more time to run (917.39 min versus 0.58 min in DrugBank). Finally, we also compared Gennius to off-the-shelf machine learning baselines Logistic Regression (LR) and Random Forest (RF), to assess the actual improvement in accuracy using the same features (see Supplementary Material Section 9 for further details on those baselines). Comparing our model with LR and RF, we observed an increase in AUROC of 31.75% and 16.73%, respectively, indicating that Gennius is superior due to its architecture: it not only uses node features but also incorporates network topological information.

**Table 2.**
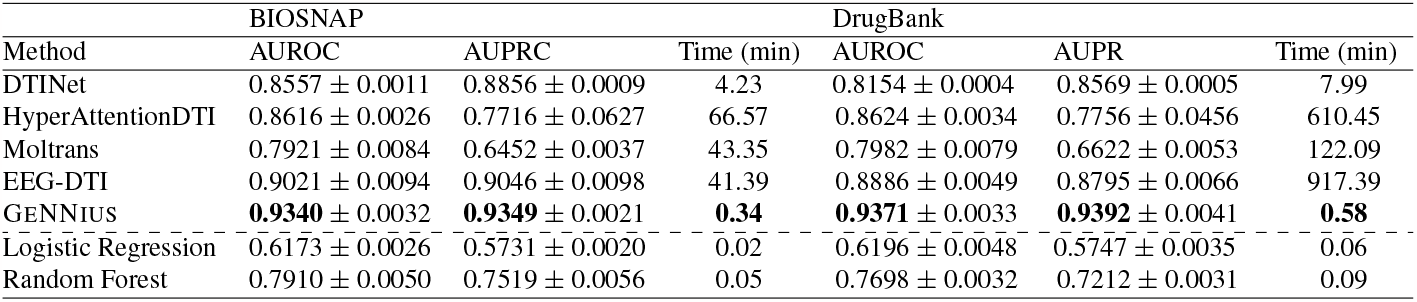
Benchmarking results of Gennius against four state-of-the-art DTI methods and two off-the-self machine learning baselines, for BIOSNAP and DrugBank datasets. Best values are highlighted in bold, excluding baseline results. All AUROC and AUPRC reported results correspond to test set, execution time correspond to train/validation/test. SOTA models were run in their default configuration, i.e., Moltrans correspond to 5 independent runs, while DTINet and EEG-DTI correspond to a 10-Fold Cross Validation, and HyperAttentionDTI to 10-times repeated 5-fold Cross-Validation.

**Fig. 2.**
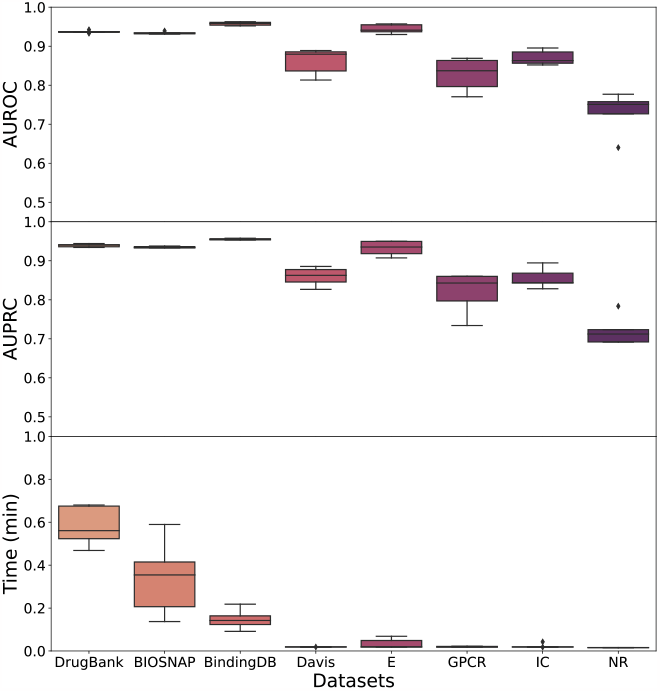
Boxplots of AUROC and AUPRC metrics on test data for five independent runs of GeNNius for the eight selected datasets. **Upper**. AUROC results. **Middle**. AUPRC results. **Lower**. Time results in minutes.

### 3.2. Gennius prediction capabilities for inferring previously unreported drug-target interactions

To analyze the capability of Gennius to detect unknown interactions, we first identified those target-protein pairs lacking an edge in one dataset (negative label) but connected in the other datasets (positive label). Then, we assessed whether Gennius was able to annotate these edges as positive. We trained the model ensuring that the edges for testing were not seen during the training process and repeated the process ten times. Further details of experiment set-up in Methods (Section 2.2.2).

The ratio of correctly predicted edges for each dataset is presented in Figure 3. When trained with large datasets, Gennius returned good prediction capabilities, detecting more than 80% of edges in almost all cases. It is worth noting that with DrugBank Gennius successfully predicted more than 90% of these edges across all runs. Further, when using Yamanishi datasets (E, GPCR, IC, and NR), Gennius returned satisfactory results, predicting 70% of DTIs on average across different runs, although with higher variability than when using large datasets. This suggests that training on a small dataset hinders the inference of new interactions, as the random choice of edges for training has larger impact on the predictive power in these cases. We note that the observed outliers could be due to a non-informative random selection of training edges. Finally, the Davis dataset yielded significantly worst results than the other datasets. At first sight, this behavior could be due to the origin of the Davis dataset, as it is generated from affinity experiments. However, BindingDB, which is also generated from affinity data, does not yield such low performance. Hence, this may indicate that the problem comes from the significant difference in the topology of Davis versus all the other datasets. Davis is the only dataset formed as a uniquely connected network, while other datasets have more than one connected component. It also presents significantly different average degree values in drug nodes (see Table 1).

**Fig. 3.**
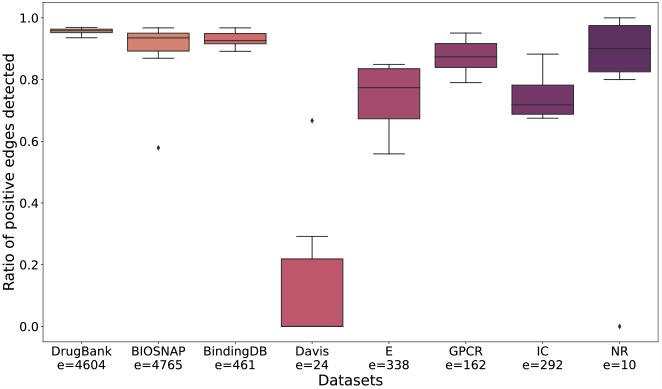
Boxplot of the ratio of correctly identified positive edges in 10 independent runs trained and tested on the same dataset. Note that *e* is the number of edges to be evaluated.

### 3.3. Gennius generalization capabilities

We evaluated Gennius performance when training and testing on different datasets. In order to ensure that there is no data leakage that might oversimplify the prediction task, DTIs that were common to train and test datasets were discarded prior to applying the model (see Methods, Section 2.2.3, for a more detailed description of the set-up).

AUROC results are presented in Figure 4 (obtained AUPRC results are similar, see Supplementary Material Section 10), where each entry of the heatmap shows the performance of Gennius on the row dataset when trained on the column dataset. The reported values correspond to 25 runs, where statistical deviation in AUROC and AUPRC arise from the random selection of edges.

**Fig. 4.**
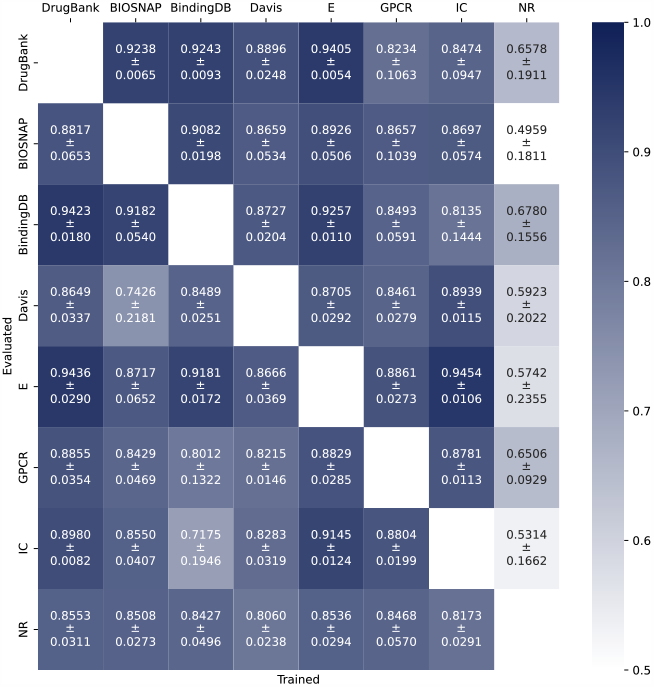
Performance of Gennius in terms of AUROC when training in one dataset (column) and testing in other (row). Train datasets do not contain positive edges that appear in the test dataset.

In general terms, Gennius returned compelling results in its generalization capabilities; however, there was a strong dependence on the training dataset. Gennius reported the best generalization capabilities when trained on larger datasets, such as DrugBank, BIOSNAP, and E. On the other hand, when the model is trained on the smallest dataset, NR, it cannot generalize, resulting in lower AUROC/AUPRC values compared to others (whiter colors in the NR column). Additionally, despite the Davis dataset being similar in size to other Yamanishi datasets, it returned the second-to-worst results for both training and testing. As mentioned previously, Davis’ topology is different from the rest of the networks. In addition, Davis and BindingDB, unlike other datasets, come from affinity experiments. However, the latter seems to perform similarly, albeit slightly worse, than DrugBank when used for training.

We also found that, for smaller networks, our method obtains better results when trained on large datasets and tested on smaller ones compared to when trained and tested on the same small dataset (see results Section 3.1). For instance, Gennius obtained an AUROC of 0.86 when trained on DrugBank and tested on NR (lower left corner of heatmap), while it achieved an AUROC of 0.73, using NR for training and testing. This suggests that training on large networks helps the model learn and generalize to unseen and smaller datasets.

In addition, to assess how much these results depend on the node features, we compared them with a random forest model that has no information on network topology. RF showed incapability to generalize, contrasting with the results obtained when training and testing on the same dataset (Table 2). The presented results indicate that Gennius is capable of generalizing by employing both features and the network’s topology without overfitting to the training network (see Supplementary Material Section 10).

Finally, we confirmed that Gennius’s generalization capability is not dependent on the embedding dimension by experimenting with different values (see Supplementary Material Section 11).

### 3.4. Gennius encoder preserves biological information in edges and diffuses it in nodes

To qualitatively interpret the generated embeddings by Gennius’s GNN encoder, we computed the t-SNE of both the input features and the computed embeddings for all nodes and DTI edges. We focused on the DrugBank dataset, since as shown in previous sections, it reports one of the best AUROC/AUPRC results (Sections 3.1 and 3.3) and yields one of the lowest variability during DTI evaluation (Section 3.2). We aimed at shedding some light on whether the embeddings generated by Gennius carry meaningful biological information beyond the ability to uncover new DTIs.

Firstly, we observe that the edge space with the input features contains information about drug chemical categories and protein families (see Supplementary Material Section 12 Figures 8a,8b,8c). Using the generated embeddings instead, we observe that despite the new shapes in the t-SNE, the biological information is conserved after graph convolutions, i.e., we can still distinguish groups by drug chemical classification but especially by protein families (see Supplementary Material Section 12 Figures 8d, 8e, 8f).

Secondly, we analyzed the nodes, and found that node input features contain almost no information about protein families, i.e., nodes do not form groups by protein families or by sub-classification of enzymes, conversely to drug nodes grouped by chemical categories (see Supplementary Material Section 12 Figures 9a, 9b, 9c). The next emerging question we wanted to answer is whether the network diffuses edge biological information during encoding such that the embedding of protein nodes reflects it. We found that the grouping of drug nodes concerning their chemical classification spread after applying the encoder; this is an awaited result, as we desire drugs in a DTI prediction model to be promiscuous (Supplementary Material Section 12 Figure 9d). However, protein node embeddings displayed better identifiable groups than before (Supplementary Material Section 12 Figure 9e). Protein families, such as membrane receptors (orange) and ion channels (violet), revealed some grouping at the top of the figure, despite not forming evident groups. Moreover, enzymes now gather in separate groups across the embedding space and, further, upon its annotation, we found a more clear grouping, e.g., kinases (fuchsia) formed a small group on the right of the t-SNE (Supplementary Material Section 12 Figure 9f).

Ultimately, the encoder maintains biological information in edge space while spreading biological information through nodes, such as protein family classification in protein nodes and sub-classification of enzymes.

Additionally, we visualized the embedding space of negative and positive edges, adding those DTI used for validation in Section 3.2. The edges embedding space differentiates areas of positive and negative edges, vastly facilitating the task to the NN-classifier (Supplementary Section 12 Figure 10).

## 4. Conclusions

We introduced a novel Drug-Target Interaction (DTI) model, termed Gennius, composed of a GNN encoder followed by an NN-edge classifier. Gennius outperformed state-ofthe-art models in terms of AUROC and AUPRC while being several orders of magnitude faster. Further, we showed that the generalization capabilities of Gennius and demonstrated its ability to infer previously unreported drug-target interactions. In addition, we showed that Gennius GNN encoder exploits both node features and graph topology to maintain biological information in edge space while spreading biological information through nodes. Ultimately, Gennius ‘s ability to generalize and predict novel DTIs reveals its suitability for drug repurposing. Additionally, its remarkable speed is key in its usability as it enables fast validation of multiple drug-target pairs.

## ACKNOWLEDGEMENTS

This work was supported by the following grants: DoD of the US - CDMR Programs [W81XWH-20-1-0262], Ramon y Cajal contracts [MCIN/AEI RYC2021-033127-I] [RYC2019-028578-I], DeepCTC [MCIN/AEI TED2021-131300B-I00], Gipuzkoa Fellows [2022-FELL-000003-01], the Spanish MCIN (PID2021-126718OA-I00), Fulbright Predoctoral Research Program [PS00342367], and FEDER/MCIN - AEI (PID2020-116344-RB-100/MCIN/AEI/10.13039/501100011033).

## Bibliography

1. Rita Santos, Oleg Ursu, Anna Gaulton, A. Patrícia Bento, Ramesh S. Donadi, Cristian G. Bologa, Anneli Karlsson, Bissan Al-Lazikani, Anne Hersey, Tudor I. Oprea, and John P. Overington. A comprehensive map of molecular drug targets. Nature Reviews Drug Discovery, 16(1):19–34, 2017. doi: 10.1038/nrd.2016.230.

2. Brittany M Duggan, Daniel M Marko, Raveen Muzaffar, Darryl Y Chan, and Jonathan D Schertzer. Kinase inhibitors for cancer alter metabolism, blood glucose, and insulin. J Endocrinol, 256(2), Feb 2023. ISSN 1479-6805 (Electronic); 0022–0795 (Linking). doi: 10.1530/JOE-22-0212.

3. Joseph A. DiMasi, Henry G. Grabowski, and Ronald W. Hansen. Innovation in the pharmaceutical industry: New estimates of r&d costs. Journal of Health Economics, 47:20–33, 2016. ISSN 0167-6296. doi: https://doi.org/10.1016/j.jhealeco.2016.01.012.

4. Sudeep Pushpakom, Francesco Iorio, Patrick A. Eyers, K. Jane Escott, Shirley Hopper, Andrew Wells, Andrew Doig, Tim Guilliams, Joanna Latimer, Christine McNamee, Alan Norris, Philippe Sanseau, David Cavalla, and Munir Pirmohamed. Drug repurposing: progress, challenges and recommendations. Nature Reviews Drug Discovery, 18(1):41–58, 2019. doi: 10.1038/nrd.2018.168.

5. Qichang Zhao, Haochen Zhao, Kai Zheng, and Jianxin Wang. HyperAttentionDTI: improving drug–protein interaction prediction by sequence-based deep learning with attention mechanism. Bioinformatics, 38(3):655–662, 10 2021.

6. Niraj Verma, Xingming Qu, Francesco Trozzi, Mohamed Elsaied, Nischal Karki, Yunwen Tao, Brian Zoltowski, Eric C. Larson, and Elfi Kraka. Ssnet: A deep learning approach for protein-ligand interaction prediction. International Journal of Molecular Sciences, 2021. doi: 10.3390/ijms22031392.

7. John Jumper, Richard Evans, Alexander Pritzel, Tim Green, Michael Figurnov, Olaf Ronneberger, Kathryn Tunyasuvunakool, Russ Bates, Augustin Žídek, Anna Potapenko, Alex Bridgland, Clemens Meyer, Simon A. A. Kohl, Andrew J. Ballard, Andrew Cowie, Bernardino Romera-Paredes, Stanislav Nikolov, Rishub Jain, Jonas Adler, Trevor Back, Stig Petersen, David Reiman, Ellen Clancy, Michal Zielinski, Martin Steinegger, Michalina Pacholska, Tamas Berghammer, Sebastian Bodenstein, David Silver, Oriol Vinyals, Andrew W. Senior, Koray Kavukcuoglu, Pushmeet Kohli, and Demis Hassabis. Highly accurate protein structure prediction with alphafold. Nature, 596(7873):583–589, 2021.

8. Mihaly Varadi, Stephen Anyango, Mandar Deshpande, Sreenath Nair, Cindy Natassia, Galabina Yordanova, David Yuan, Oana Stroe, Gemma Wood, Agata Laydon, Augustin Žídek, Tim Green, Kathryn Tunyasuvunakool, Stig Petersen, John Jumper, Ellen Clancy, Richard Green, Ankur Vora, Mira Lutfi, Michael Figurnov, Andrew Cowie, Nicole Hobbs, Pushmeet Kohli, Gerard Kleywegt, Ewan Birney, Demis Hassabis, and Sameer Velankar. AlphaFold Protein Structure Database: massively expanding the structural coverage of protein-sequence space with high-accuracy models. Nucleic Acids Research, 50(D1): D439–D444, 11 2021.

9. Kexin Huang, Cao Xiao, Lucas M Glass, and Jimeng Sun. MolTrans: Molecular Interaction Transformer for drug–target interaction prediction. Bioinformatics, 37(6):830–836, 10 2020.

10. Yunan Luo, Xinbin Zhao, Jingtian Zhou, Jinglin Yang, Yanqing Zhang, Wenhua Kuang, Jian Peng, Ligong Chen, and Jianyang Zeng. A network integration approach for drug-target interaction prediction and computational drug repositioning from heterogeneous information. Nat Commun, 8(1):573, 2017.

11. Jiajie Peng, Yuxian Wang, Jiaojiao Guan, Jingyi Li, Ruijiang Han, Jianye Hao, Zhongyu Wei, and Xuequn Shang. An end-to-end heterogeneous graph representation learning-based framework for drug–target interaction prediction. Briefings in Bioinformatics, 22(5), 2021.

12. Cong Xu, Huiling Huang, Xiaoting Ying, Jianliang Gao, Zhao Li, Peng Zhang, Jie Xiao, Jiarun Zhang, and Jiangjian Luo. Hgnn: Hierarchical graph neural network for predicting the classification of price-limit-hitting stocks. Information Sciences, 607:783–798, 2022. ISSN 0020-0255. doi: https://doi.org/10.1016/j.ins.2022.06.010.

13. Zhiqiang Que, Marcus Loo, and Wayne Luk. Reconfigurable acceleration of graph neural networks for jet identification in particle physics. In 2022 IEEE 4th International Conference on Artificial Intelligence Circuits and Systems (AICAS), 2022. doi: 10.1109/AICAS54282.2022.9869941.

14. Hai-Cheng Yi, Zhu-Hong You, De-Shuang Huang, and Chee Keong Kwoh. Graph representation learning in bioinformatics: trends, methods and applications. Briefings in Bioinformatics, 23(1), 09 2021.

15. William L. Hamilton. Graph representation learning. Synthesis Lectures on Artificial Intelligence and Machine Learning, 14(3):1–159, 2020.

16. Aditya Grover. node2vec: Scalable feature learning for networks, 2016.

17. Matthias Fey and Jan E. Lenssen. Fast graph representation learning with PyTorch Geometric. In RLGM Workshop at ICLR, 2019.

18. William L. Hamilton, Rex Ying, and Jure Leskovec. Inductive representation learning on large graphs. CoRR, abs/1706.02216, 2017.

19. Lingfei Wu, Peng Cui, Jian Pei, and Liang Zhao. Graph Neural Networks: Foundations, Frontiers, and Applications. Springer Singapore, Singapore, 2022.

20. Diederik P. Kingma. Adam: A method for stochastic optimization, 2017.

21. Adam Paszke, Sam Gross, Francisco Massa, Adam Lerer, James Bradbury, Gregory Chanan, Trevor Killeen, Zeming Lin, Natalia Gimelshein, Luca Antiga, Alban Desmaison,Andreas Köpf, Edward Yang, Zach DeVito, Martin Raison, Alykhan Tejani, Sasank Chilamkurthy, Benoit Steiner, Lu Fang, Junjie Bai, and Soumith Chintala. PyTorch: An Imperative Style, High-Performance Deep Learning Library. Curran Associates Inc., Red Hook, NY, USA, 2019.

22. Stefan Wager, Sida Wang, and Percy Liang. Dropout training as adaptive regularization, 2013.

23. Greg Landrum, Paolo Tosco, Brian Kelley, Gedeck Sriniker, and Gedeck. Rdkit: Open-source cheminformatics. 2022.

24. David S Wishart, Craig Knox, An Chi Guo, Savita Shrivastava, Murtaza Hassanali, Paul Stothard, Zhan Chang, and Jennifer Woolsey. Drugbank: a comprehensive resource for in silico drug discovery and exploration. Nucleic acids research, 2006.

25. Sagar Maheshwari Marinka Zitnik, Rok Sosič and Jure Leskovec. BioSNAP Datasets: Stan-ford biomedical network dataset collection, 2018.

26. Stanford-SNAP-Group. Miner: Gigascale multimodal biological network. GitHub Repository, 2017.

27. Tiqing Liu, Yuhmei Lin, Xin Wen, Robert N Jorissen, and Michael K Gilson. Bindingdb: a web-accessible database of experimentally determined protein–ligand binding affinities. Nucleic acids research, 35(suppl_1)D198–D201, 2007.

28. Kexin Huang, Tianfan Fu, Wenhao Gao, Yue Zhao, Yusuf Roohani, Jure Leskovec, Connor W. Coley, Cao Xiao, Jimeng Sun, and Marinka Zitnik. Therapeutics data commons: Machine learning datasets and tasks for drug discovery and development, 2021.

29. Mindy I Davis, Jeremy P Hunt, Sanna Herrgard, Pietro Ciceri, Lisa M Wodicka, Gabriel Pallares, Michael Hocker, Daniel K Treiber, and Patrick P Zarrinkar. Comprehensive analysis of kinase inhibitor selectivity. Nature biotechnology, 29(11):1046–1051, 2011.

30. Yoshihiro Yamanishi, Michihiro Araki, Alex Gutteridge, Wataru Honda, and Minoru Kanehisa. Prediction of drug–target interaction networks from the integration of chemical and genomic spaces. Bioinformatics, 2008.

31. Nansu Zong, Rachael Sze Nga Wong, Yue Yu, Andrew Wen, Ming Huang, and Ning Li. Drug–target prediction utilizing heterogeneous bio-linked network embeddings. Briefings in Bioinformatics, 22(1):568, 12 2019. doi: 10.1093/bib/bbz147.

32. Junjun Zhang and Minzhu Xie. Graph regularized non-negative matrix factorization with prior knowledge consistency constraint for drug–target interactions prediction. BMC Bioinformatics, 23(1):564, 2022. doi: 10.1186/s12859-022-05119-6.

33. Minoru Kanehisa, Susumu Goto, Masahiro Hattori, Kiyoko F Aoki-Kinoshita, Masumi Itoh, Shuichi Kawashima, Toshiaki Katayama, Michihiro Araki, and Mika Hirakawa. From genomics to chemical genomics: new developments in kegg. Nucleic Acids Res, 34(Database issue):D354–7, Jan 2006.

34. Ida Schomburg, Antje Chang, Christian Ebeling, Marion Gremse, Christian Heldt, Gregor Huhn, and Dietmar Schomburg. Brenda, the enzyme database: updates and major new developments. Nucleic Acids Res, 32(Database issue):D431, Jan 2004.

35. Stefan Günther, Michael Kuhn, Mathias Dunkel, Monica Campillos, Christian Senger, Evangelia Petsalaki, Jessica Ahmed, Eduardo Garcia Urdiales, Andreas Gewiess, Lars Juhl Jensen, Reinhard Schneider, Roman Skoblo, Robert B Russell, Philip E Bourne, Peer Bork, and Robert Preissner. Supertarget and matador: resources for exploring drug-target relationships. Nucleic Acids Res, 36(Database issue):D919, Jan 2008.

36. Anna Gaulton, Anne Hersey, Michał Nowotka, A. Patrícia Bento, Jon Chambers, David Mendez, Prudence Mutowo, Francis Atkinson, Louisa J. Bellis, Elena Cibrián-Uhalte, Mark Davies, Nathan Dedman, Anneli Karlsson, María Paula Magariños, John P. Overington, George Papadatos, Ines Smit, and Andrew R. Leach. The ChEMBL database in 2017. Nucleic Acids Research, 45(D1):D945–D954, 11 2016. ISSN 0305-1048. doi: 10.1093/nar/gkw1074.

37. Yannick Djoumbou Feunang, Roman Eisner, Craig Knox, Leonid Chepelev, Janna Hastings, Gareth Owen, Eoin Fahy, Christoph Steinbeck, Shankar Subramanian, Evan Bolton, Russell Greiner, and David S. Wishart. Classyfire: automated chemical classification with a comprehensive, computable taxonomy. Journal of Cheminformatics, 8(1):61, 2016.

