## Supplementary Material for "GeNNius: An ultrafast drug-target interaction inference method based on graph neural networks"

### 1. Hyperparameter tuning for GeNNius's GNN encoder

We performed a grid search to asses which hyperparameters are more effective for the DTI prediction task. The grid search was performed using the DrugBank dataset, for being the largest dataset available, under the idea that it may be easier for the model to generalize. The grid search tested several hyperparameters, repeating ten times each hyperparameter configuration. In all cases the dataset was randomly split into train, validation and test sets with a 70/10/20 ratio. For selecting the best-performing architecture, we used the validation set to prevent overfitting to the test set, the latter used to report results in the main text. The script used to perform the hyperparameter tuning is provided in the GitHub repository (`https://github.com/ubioinformat/GeNNius`), as well as a CSV file with the obtained results.
We evaluated the following hyperparameters:

- Type of layer: SAGEConv, GraphConv, GATConv, Tranformerconv. For further information about the layers and their implementation visit the PyG webpage (`https://pytorch-geometric.readthedocs.io/en/latest/modules/nn.html`). Note even if there are several layers implemented in PyG, not all can be used in the scenario of an heterogeneous bipartite network.
- Number of layers: from one to five, as with more than 4 layers the AUROC/AUPRC decreased considerably due to the over smoothing problem.
- Embedding dimension: from 6 to 24.
- Dropout rate: 0, 0.2, and 0.5.
- Number of heads for those models with attention (GATConv, Transformerconv): 1, 2, 4.
- Aggregators for SAGEConv and GraphConv layers: sum, mean, max.
- Activation functions: ReLu and Tanh.

The best model (i.e., hyperparameter configuration) was selected as the one with the highest average AUROC (and AUPRC) on the validation set across the ten independent runs. While the variation of some hyperparameters, such as the embedding dimension, did not affect AUROC results, others did, such as the aggregator function of the convolutional layer and the dropout rate. Figures 1a, and 1b show the obtained results as a function of the aggregator function and the dropout rate, respectively. The SUM aggregator function provided the highest AUROC values, and the dropout rate of 0.2 enhances the model's learning compared to higher values (0.5) and to no dropout at all.

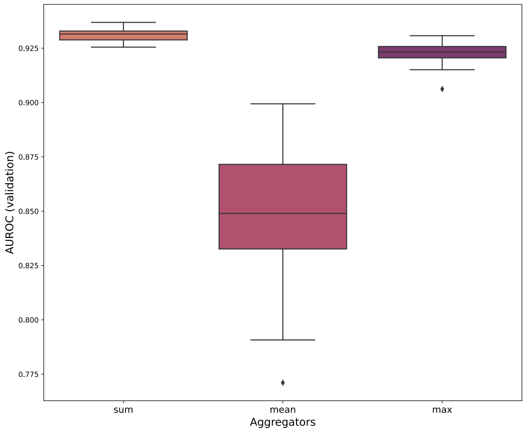


**(a)** Boxplots of the AUROC as a function of the aggregator function.

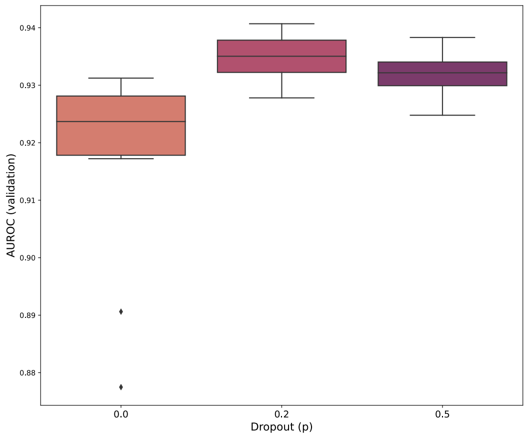


**(b)** Boxplots of the AUROC as a function of the dropout rate.

**Fig. 1.** Boxplot of the AUROC values obtained on the validation set with GeNNius selected architecture but varing two different hyperparameters. (a) AUROC boxplot as a function of the aggregator function., and (b) Boxplots of the AUROC as a function of the dropout rate.

The selected hyperparameters for GeNNius's GNN architecture are summarized in Table 3.

**Table 3.** Summary of GeNNius hyperparameters.

| Parameter type | Value |
|---|---|
| Layer type | SAGEConv |
| Number of hidden layer | 4 |
| Activation function | Tanh |
| Aggregation type | Sum |
| Embedding dimension | 17 |
| Learning rate | 0.01 |
| Dropout | 0.2 |

### 2. Evaluation metrics

To assess the performance of the models in the classification task, we used the area under the ROC (Receiver Operating Characteristic) curve (AUROC), as well as the area under the precision-recall curve (AUPRC), both widely used for evaluating DTI prediction models. The ROC curve depicts the false positive rate (FPR) versus the true positive rate (TPR), defined as:

$$TPR = \frac{TP}{TP+FN}; \quad FPR = \frac{FP}{FP+TN}, \tag{5}$$

where TP, FN, and FP stand for true positives, false negatives, and false positives, respectively. The precision-recall (PR) curve plots instead the precision (P) versus the recall (R), defined as follows:

$$P = \frac{TP}{TP+FP}; \quad R = TPR = \frac{TP}{TP+FN}. \tag{6}$$

Recall that for a given input corresponding to the information of a drug and protein node, the classifier outputs a number between 0 and 1, indicating the probability of existence of and edge between the two nodes (i.e., a positive outcome). By modifying the threshold for which a decision between negative and positive outcomes is made, we can generate the ROC and PR curves.

The AUROC and AUPRC are then given by the area under the ROC or PR curves, respectively, and ranges between 0 and 1, with 0.5 corresponding to a random classifier and 1 to a perfect one. This value helps to evaluate the reliability and confidence of the models. Intuitively, positive (negative) inputs should produce high (low) probabilities. Sorting the inputs by their probability should therefore result in positive samples appearing before negative ones. In other words, a high (low) threshold should produce few FPs (FNs). Hence, the considered AUROC and AUPRC metrics show how well the model separates both classes.

### 3. Node Features correlation

Correlation maps of drug node features for DrugBank dataset in Figure 2a. For completeness, Figure 2b presents the correlation matrix of protein features, also for DrugBank.

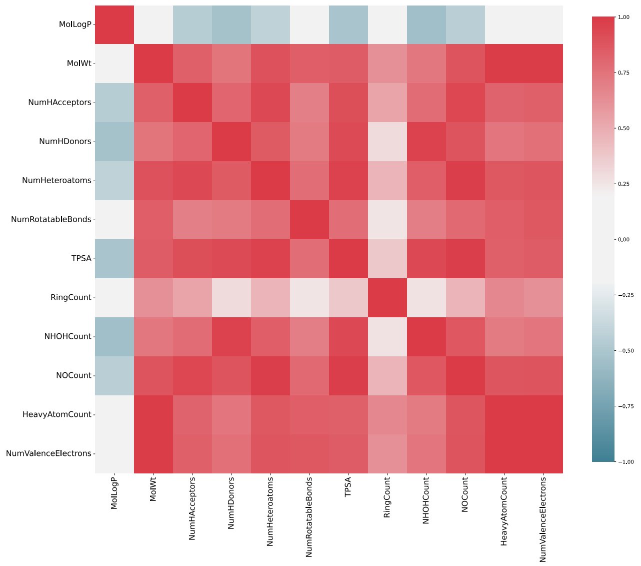


**(a)** Drug features correlation matrix

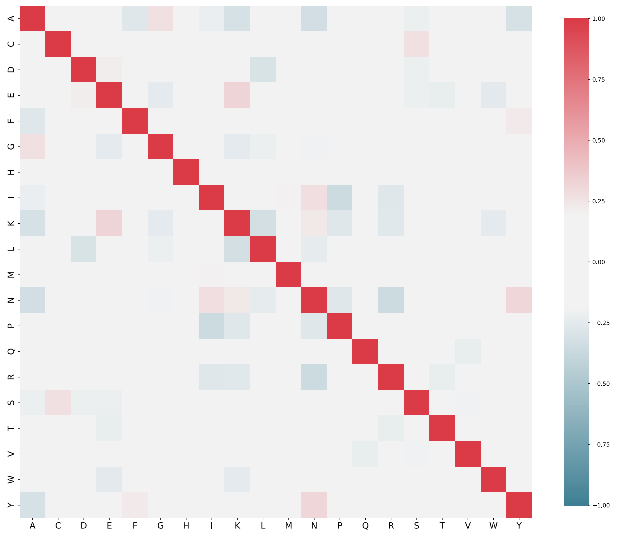


**(b)** Protein features correlation matrix

**Fig. 2.** Correlation of node features in DrugBank. (a) Drug nodes feature correlation and (b) protein node features correlation.

### 4. Using pre-computed protein embeddings

To test other protein features that could be more standard in other tasks or informative to the model, we trained and tested the model using pre-computed embeddings of protein sequences, specifically embeddings retrieved from the UniProt database, corresponding to those generated using the ProtT5 model (https://www.uniprot.org/help/embeddings).
The use of those features, with the same architecture selected with GeNNius, did not increase considerably the resulting AUROC (see Table 4). Further, it seems that for smaller models such as NR the model is overfitting; for 2 out of 5 runs, NR returned 1.000 in AUROC. Moreover, in terms of generalization seemed to hamper the model. Firstly, smaller networks such as NR pre-computed embeddings were not available at Uniprot, leading to a network with not enough edges for training. Secondly, the generalization results for the rest of the available datasets were worse for some datasets (see heatmap in Figure 3). These results validate the use of amino acid ratio for the initial protein features, being also faster to generate, and available for all proteins.

**Table 4.** Results with protein pre-computed embedding as protein node features for the selected datasets.

| Dataset | AUROC | AUPRC | Time |
|---|---|---|---|
| DrugBank | $0.9304 \pm 0.0072$ | $0.9329 \pm 0.0070$ | $0.15 \pm 0.09$ |
| BIOSNAP | $0.9263 \pm 0.0061$ | $0.9306 \pm 0.0070$ | $0.10 \pm 0.01$ |
| BindingDB | $0.9475 \pm 0.0094$ | $0.9445 \pm 0.0110$ | $0.04 \pm 0.02$ |
| Davis | $0.6325 \pm 0.0561$ | $0.6554 \pm 0.0806$ | $0.02 \pm 0.00$ |
| E | $0.9382 \pm 0.0060$ | $0.9278 \pm 0.0080$ | $0.03 \pm 0.01$ |
| GPCR | $0.8212 \pm 0.0163$ | $0.8257 \pm 0.0240$ | $0.02 \pm 0.00$ |
| IC | $0.8609 \pm 0.0107$ | $0.8506 \pm 0.0145$ | $0.02 \pm 0.01$ |
| NR | $0.8625 \pm 0.1556$ | $0.8767 \pm 0.1323$ | $0.02 \pm 0.00$ |

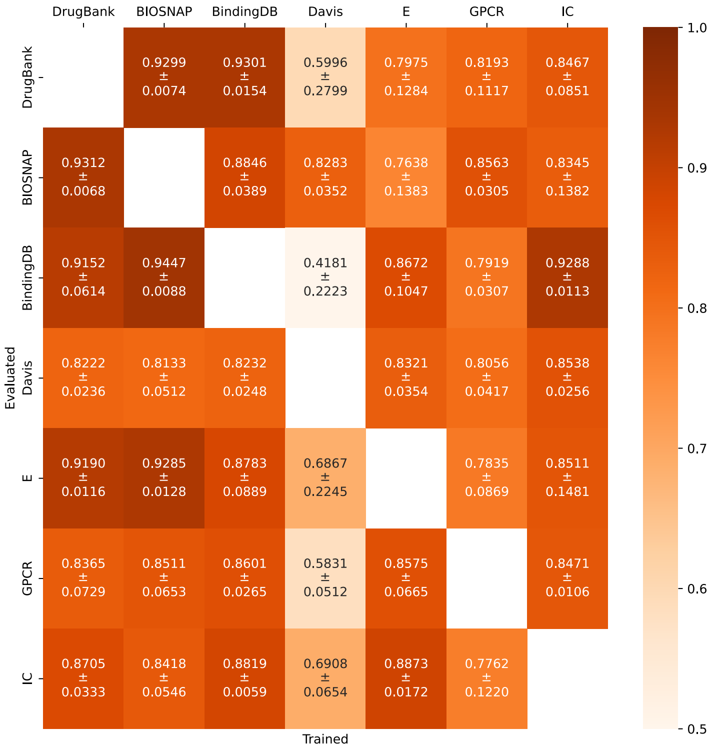


**Fig. 3.** Performance of GeNNius in terms of AUROC using pre-computed embeddings as initial features ($d = 17$ for GNN encoder) when training in one dataset (column) and testing in other (row). Train datasets do not contain positive edges that appear in the testing dataset. Set-up similar to that explained in Main Section 2.2.3.

### 5. Additional datasets information

Table 5 specifies, for each considered dataset, the number of drugs, proteins, nodes and edges, originally and after dropping those nodes for which features could not be generated.

**Table 5.** Datasets statistics originally and in brackets lost nodes or edges.

| | | | | | Yamanishi | | | |
|---|---|---|---|---|---|---|---|---|
| | DrugBank | BIOSNAP | BindingDB | Davis | E | GPCR | IC | NR |
| Number of drugs | 6839 (16) | 4502 (3) | 3085 (1) | 59 (0) | 444 (0) | 222 (0) | 210 (0) | 53 (0) |
| Number of proteins | 4663 (11) | 2122 (9) | 719 (1) | 218 (2) | 660 (0) | 94 (0) | 204 (0) | 25 (0) |
| Total number of nodes | 11502 (27) | 6624 (12) | 3804 (2) | 277 (2) | 1104 (0) | 316 (0) | 414 (0) | 78 (0) |
| Total number of edges | 24243 (535) | 13879 (41) | 5942 (5) | 673 (6) | 2920 (0) | 634 (0) | 1476 (0) | 88 (2) |

### 6. Edge analysis between datasets

In order to asses the similarities and dissimilarities between datasets, we generated four different heatmaps summarizing the percentage of shared edges (Figure 4a), and the amount of edges not reported as positive in one dataset but that were found to be positive in others (Figure 4b).

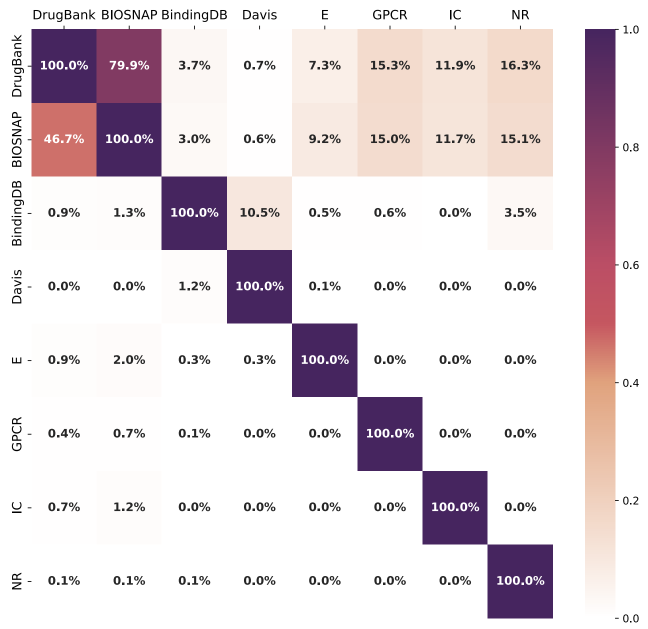


**(a)** Heatmap of repeated edges.

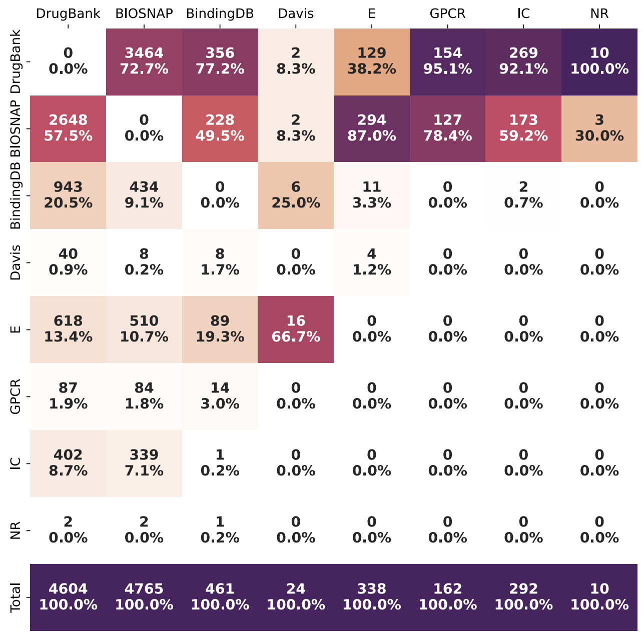


**(b)** Heatmap of negative edges.

**Fig. 4.** Datasets comparison statistics. (a) heatmap of repeated edges, where each entry represents the percentage of edges that the dataset in the column shares with the one in the row. (b) Heatmap of negative edges. Each entry represents first the number of negative edges from the dataset in the column that has been registered as positives in the one in the row, and second, the percentage of edges per dataset compared to the total number to evaluate, numbers that correspond to those in Main Section 3.2. Note that some edges may be repeated, for that reason the total number may be lower than the sum of edges per dataset.

### 7. Additional results

Table 6 provides the average and standard deviation of the AUROC and AUPRC metrics obtained by GeNNius on the different datasets (always in the test set) across five independent runs. The running time corresponds to training, validation and testing. The same results are represented in main text Figure 2.

**Table 6.** Results corresponding to main text Figure 2.

| Dataset | AUROC | AUPRC | Time (min) |
|---|---|---|---|
| DrugBank | $0.9371 \pm 0.0033$ | $0.9392 \pm 0.0041$ | $0.58 \pm 0.09$ |
| BIOSNAP | $0.9339 \pm 0.0032$ | $0.9349 \pm 0.0021$ | $0.34 \pm 0.18$ |
| BingingDB | $0.9576 \pm 0.0045$ | $0.9552 \pm 0.0019$ | $0.15 \pm 0.05$ |
| Davis | $0.8607 \pm 0.0338$ | $0.8596 \pm 0.0238$ | $0.02 \pm 0.00$ |
| E | $0.9440 \pm 0.0117$ | $0.9321 \pm 0.0190$ | $0.03 \pm 0.02$ |
| GPCR | $0.8273 \pm 0.0428$ | $0.8189 \pm 0.0540$ | $0.02 \pm 0.00$ |
| IC | $0.8704 \pm 0.0189$ | $0.8557 \pm 0.0260$ | $0.02 \pm 0.01$ |
| NR | $0.7304 \pm 0.0536$ | $0.7207 \pm 0.0378$ | $0.02 \pm 0.00$ |

### 8. Running time in CPU

GeNNius was also run without GPU to check the difference in time with respect to GPU (see main text Table 7). Reported running times correspond to train, test and validation.

**Table 7.** Time results (average and std) in minutes when running the model with CPU for all considered datasets. Presented results correspond to an average of 5 runs.

| | Time | |
|---|---|---|
| Dataset | Average | std |
| DrugBank | 0.9188 | 0.3297 |
| BIOSNAP | 0.6690 | 0.2445 |
| BindingDB | 0.1911 | 0.0845 |
| Davis | 0.0188 | 0.0001 |
| E | 0.0534 | 0.0295 |
| GPCR | 0.0188 | 0.0001 |
| IC | 0.0240 | 0.0068 |
| NR | 0.0159 | 0.0002 |

### 9. Baseline models

To assess whether the node features or the model itself (which includes information on network topology) play an essential role, i.e., if the obtained performance metrics are mostly due to the used features, we used two baseline models that use only node features for prediction: Logistic Regression (LR) and Random Forest (RF).
For hyperparameter tuning, we used the DrugBank dataset, for being the largest dataset used in this work. The models were implemented with the sklearn Python package. Both baselines take edges (concatenation of protein and drug features) as input to classify whether they are labeled as positive (interaction) or negative (not interaction). Those edges were selected by using the same function as in GENNIUS, i.e., using the *RandomLinkSplit* function by PyG. Code available in the GitHub repository.

#### Logistic Regression

Logistic regression (LR) is a statistical model used for binary classification problems, where the goal is to predict the probability of a binary outcome. It estimates the relationship between the input and the output variable, using a logistic function to transform the input into a probability value between 0 and 1. The selected hyperparameters are presented in Table 8.

**Table 8.** Summary of LR hyperparameters

| Parameter type | Value |
|---|---|
| C | 3.16 |
| Max_iter | 100 |
| Penalty | ‘l2’ |

#### Random Forest

Random forest (RF) is an ensemble model that trains multiple decision trees and combines their outputs to make predictions. Each decision tree is built using a random subset of features and training data, which helps to reduce overfitting and increase accuracy. The selected hyperparameters are presented in Table 9.

**Table 9.** Summary of RF hyperparameters

| Parameter type | Value |
|---|---|
| criterion | gini |
| max_depth | 10 |
| max_features | auto |
| min_samples_leaf | 6 |
| min_samples_split | 5 |
| n_estimators | 50 |

### 10. Additional generalization results

The generalization AUPRC results of GeNNius are shown in Figure 5. For completeness, we also tested the generalization ability when no network topology is used. The results when using the RF baseline model in terms of AUROC and AUPRC are depicted in Figures 6a and 6b, respectively. We selected the RF model as it exhibited better performance than the LR model (see main Table 2). Details on the RF model and hyperparameters are provided in Supplementary Section 9.

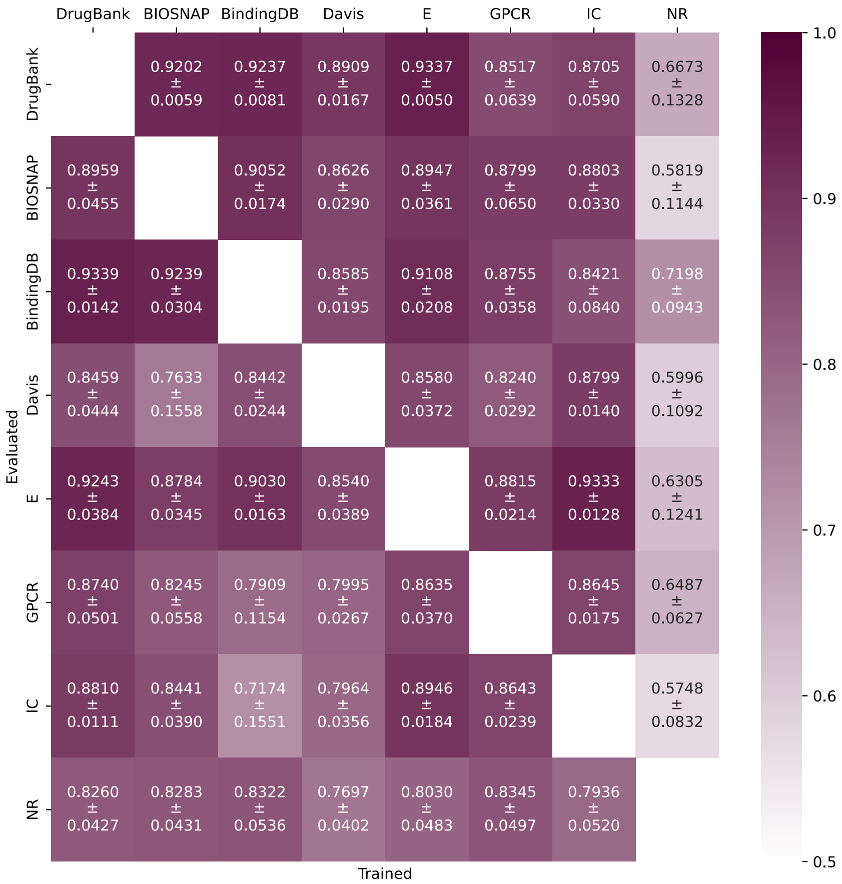


**Fig. 5.** Performance of GeNNius in terms of AUPRC when training in one dataset (column) and testing in other (row). Train datasets do not contain positive edges that appear in the testing dataset.

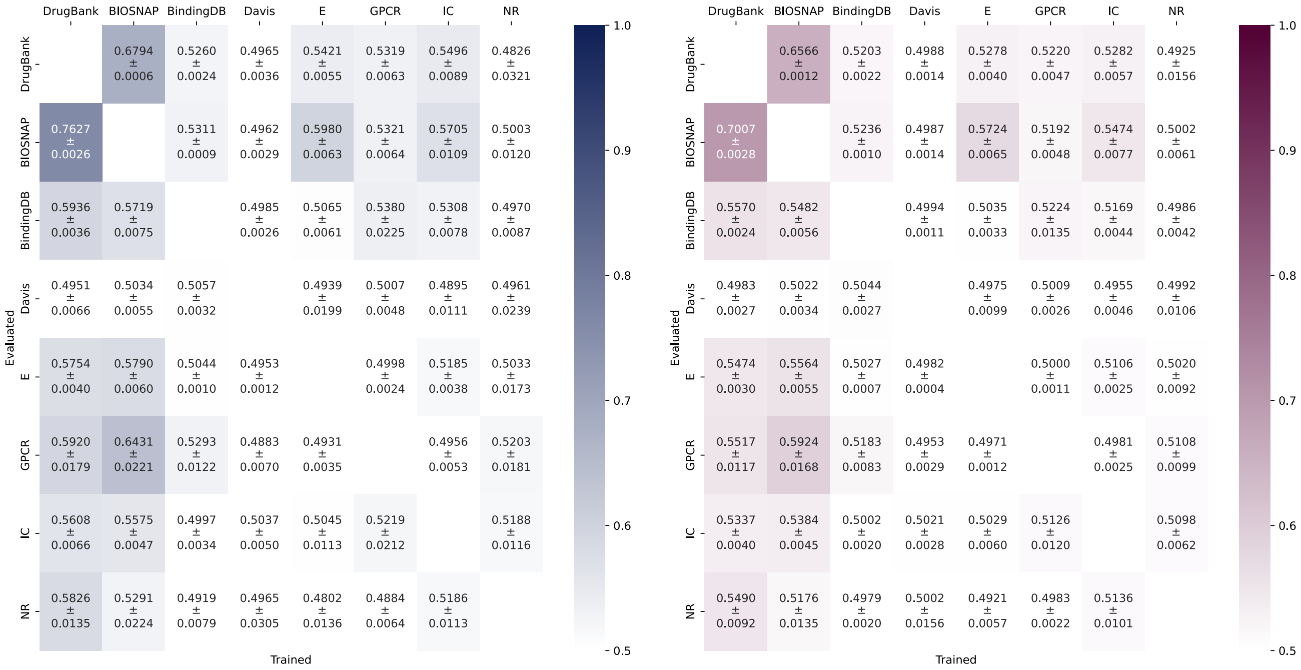


**(a)** AUROC Heatmap Baseline RF

**(b)** AUPRC Heatmap Baseline RF

**Fig. 6.** Generalization performance of Random Forest in terms of (a) AUROC and (b) AUPRC (average and std), when training in one dataset (column) and testing in other (row). Train datasets do not contain positive edges that appear in the testing dataset. Results correspond to 25 runs, see methods in main article for further details on edge selection.

### 11. Selected embedding dimension does not hamper generalization

To check whether the embedding dimension ($d = 17$) selected via grid-search with DrugBank may hamper the generalization capabilities of the model, we repeated the generalization experiment with different embedding dimensions. As explained in the main text (Section 2.2.3), we train and test the model on different datasets, for all possible combinations. For each train-test pair, we run the model 25 times.

Results are presented in Figure 7. As it can be observed, there are almost no changes in AUROC and AUPRC as a function of the embedding dimension. These results indicate that the selected $d = 17$ embedding dimension does not hamper the generalization capabilities of GENNIUS.

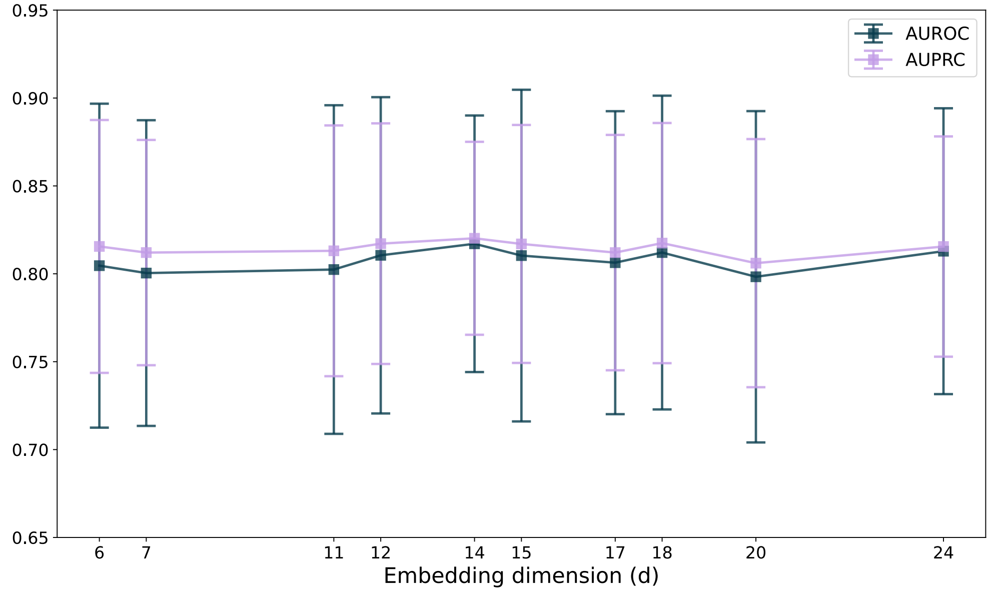


**Fig. 7.** Average and standard deviation of AUROC and AUPRC values when training and testing on different datasets, for different embedding dimensions.

### 12. Edge and node embedding analysis

We wanted to qualitatively interpret how our GENNIUS' encoder manages biological information in the network. For this, we used the DrugBank dataset; it is the largest dataset and also returned satisfactory results for all conducted experiments.
Firstly, we retrieved all edges in the network and concatenated the corresponding nodes's initial features to get a view of the possible biological information stored in the network's topology. Then, with the resulting data, we performed dimensionality reduction via t-SNE. The results indicate that the network contains information about protein families and drug classification (Figures 8a, 8e, 8c). We can clearly distinguish edges annotated by the same drug classification grouping together (Figure 8a). The same is observed for edges annotated by protein classification, such as groups of secreted enzymes (red), transporters (green), and (ion channels). These three examples of proteins show that the information on protein families in the network appears on the edges (Figure 8e). Similarly, this grouping appeared when annotating by enzyme subclassification (Figure 8c). Next, we generated the t-SNE of the embedding space of edges to inspect whether the encoder maintains the biological information in this space and we verified that, despite the new shapes in the t-SNE, the biological information is conserved after graph convolutions, i.e., we can still distinguish edge groups by drug chemical classification but especially by protein families (see Figures 8d, 8e, 8f).

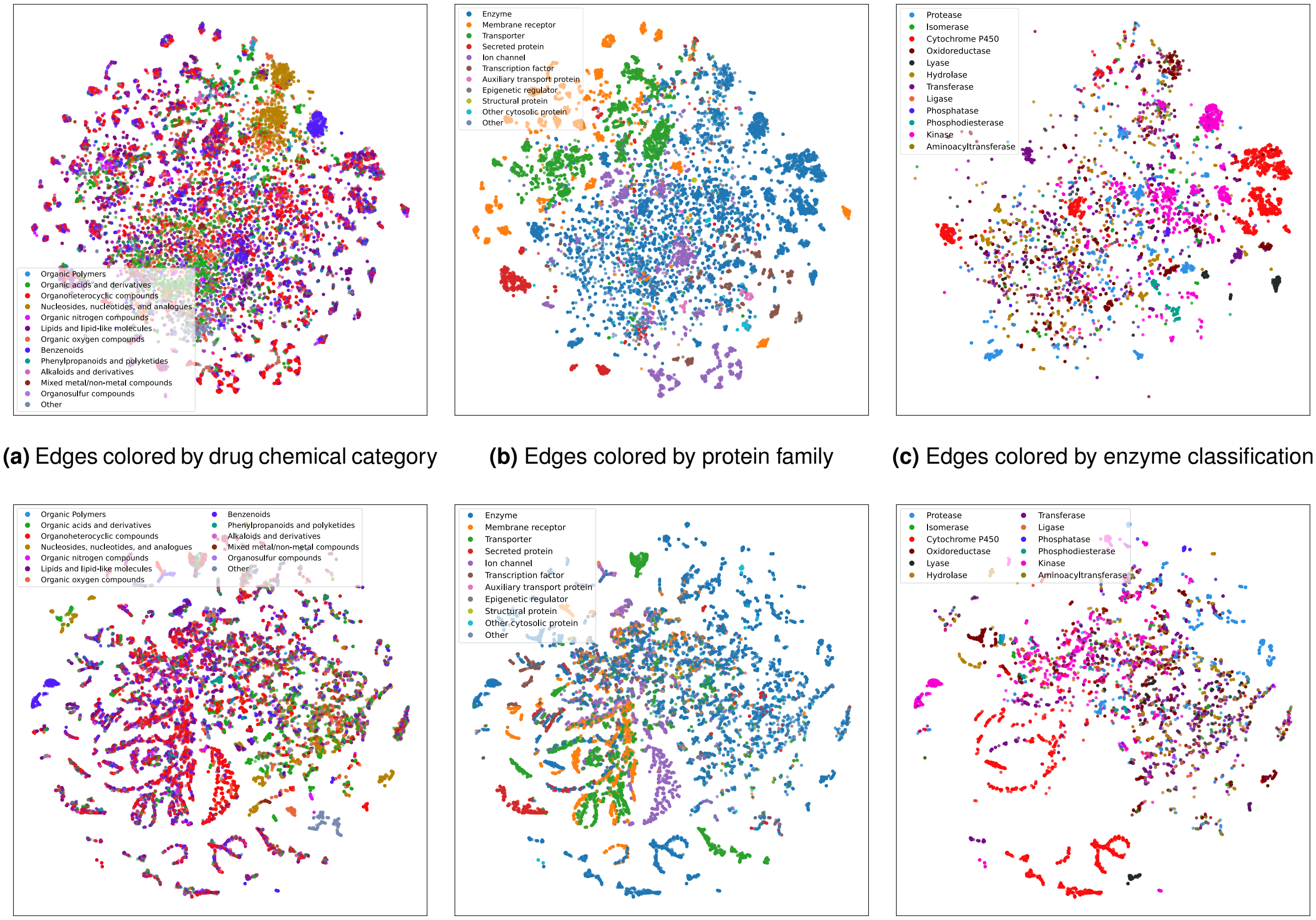


**(a)** Edges colored by drug chemical category **(b)** Edges colored by protein family **(c)** Edges colored by enzyme classification

**(d)** Edge colored by drug chemical category **(e)** Edge colored by protein family **(f)** Edge colored by enzyme classification

**Fig. 8.** t-SNE representation of the edge space. t-SNE parameters for all t-SNEs: perplexity = 30, niter= 1000, selected metric was cosine with random initialization. For drugs, removed unclassified drugs, those who belong to "Other" group and those with less than 20 appearances together. For proteins removed unclassified proteins and those with less than 10 appearances.

Next we analyzed the nodes themselves, rather than the edges. Drug features grouped in the reduced space in a manner that resembled family branches or leaves, as expected due to the high relation between molecular descriptors and drug chemical categories (Figure 9a). Protein features, conversely, showed little to no evidence of containing information about their respective families. We could only distinguish a separation for proteins corresponding to transporters and membrane receptors; for those, this is an expected behavior, as those proteins evolved to 3D structures with membrane-bound functions, including more presence of hydrophobic amino acids (attach the protein in the lipid bilayer) and less of charged amino acids (avoid repulsion) (Figure 9b). Further, enzymes are soluble proteins that catalyze chemical reactions in the cell and contain specialized active sites. Hence, it may be expected an amino acid ratio with a higher proportion of polar or charged amino acids in the active site to interact with and stabilize, which may also depend on the type of enzyme. Some slight displacement of kinases, shown in pink in Figure 9c), can be appreciated, but no strong grouping was found.

After applying GeNNius' encoder, we proceeded similar but plotting the embedded space of protein and drug nodes. We found that the grouping of drug nodes concerning their chemical classification spread after applying the encoder; this is a compelling result, as we desire drugs in a DTI prediction model to be promiscuous (Figure 9d). However, protein node embeddings displayed better identifiable groups (Figure 9e). Protein families, such as membrane receptors (orange) and ion channels (violet), revealed some grouping at the top of the Figure, despite not forming evident groups. Further, enzymes now gather in separate groups across the embedding space and, further, upon its annotation, we found a more clear group; kinases (fuchsia) formed a small group on the right of the plot; relatively similar, cytochrome P450 (red) appeared in a small group at the top; proteases (light blue) form a small group in the middle, and some oxidoreductases (brown) grouped at the left of the plot (Figure 9f).

Ultimately, the encoder maintains biological information in edge space while spreading biological information through nodes, such as protein family classification in protein nodes and sub-classification of enzymes.

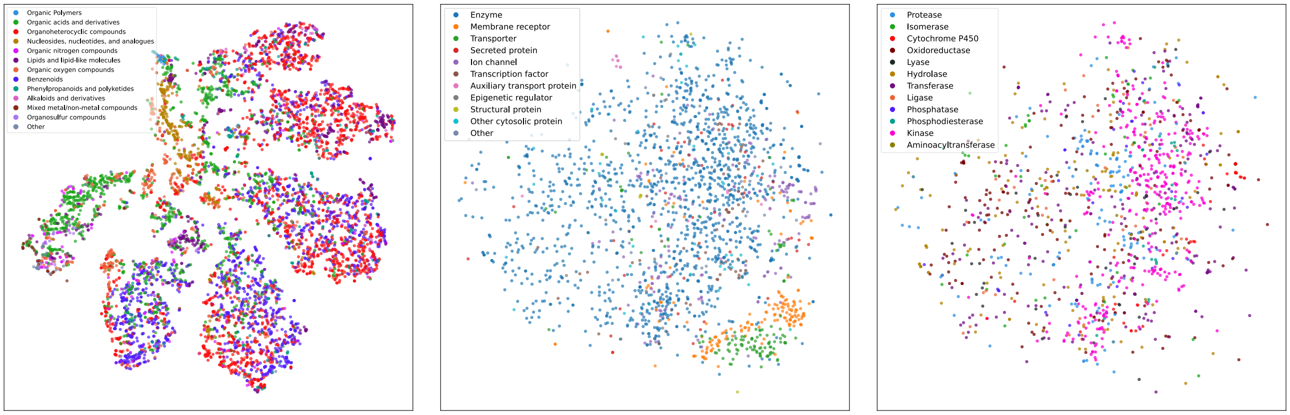


**(a)** Drug Features colored by chemical categories **(b)** Protein features colored by protein family **(c)** Protein features colored by enzyme classification

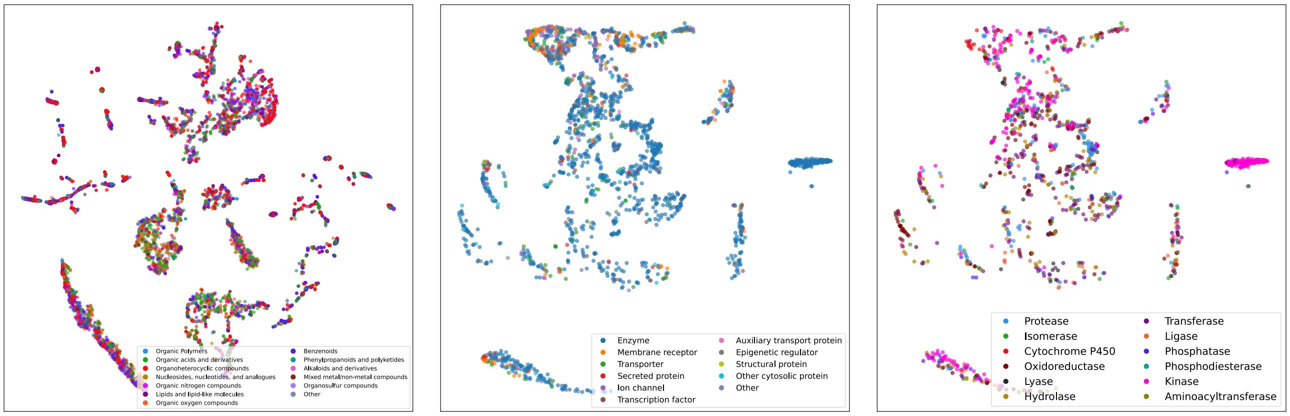


**(d)** Drug embedding colored by chemical categories **(e)** Protein embedding colored by protein family **(f)** Protein embedding colored by enzyme classification

**Fig. 9.** t-SNE representation of the nodes space. t-SNE parameters: **(a,b,c)** perplexitiy = 45, niter= 850, using euclidean metric with random initialization. t-SNE parameters; **(d,e,f)** perplexitiy = 30, number of iterations= 1000, using cosine metric and random initialization. For drugs, removed unclassified drugs, those who belong to "Other" group and those with less than 20 appearances together. For proteins removed unclassified proteins and those with less than 10 appearances.

Finally, we visualized the embedding space of negative and positive edges, adding those DTI used for validation in Main Section 3.2. This section reports results for the DrugBank dataset.
The edges in the embedding space differentiate areas of positive and negative edges, vastly facilitating the task to the NN-classifier. In both plots of Figure 10, the edges for training are represented in light colors, labeled positive (red) and negatively (blue) depending if they define a DTI or nor, respectively. Negatives were selected randomly. Then, edges used for validation in Main Section 3.2 are depicted in dark colors. Recall that these are negative edges that were found to be positive in other datasets. In Figure 10-Left, the output probability of edges to be positive has been binarized using a threshold of 0.5, i.e., edges with a probability greater (smaller) than 0.5 are predicted as positive (negative). In Figure 10-Right, the output probability of edges has been plotted in greyscale.

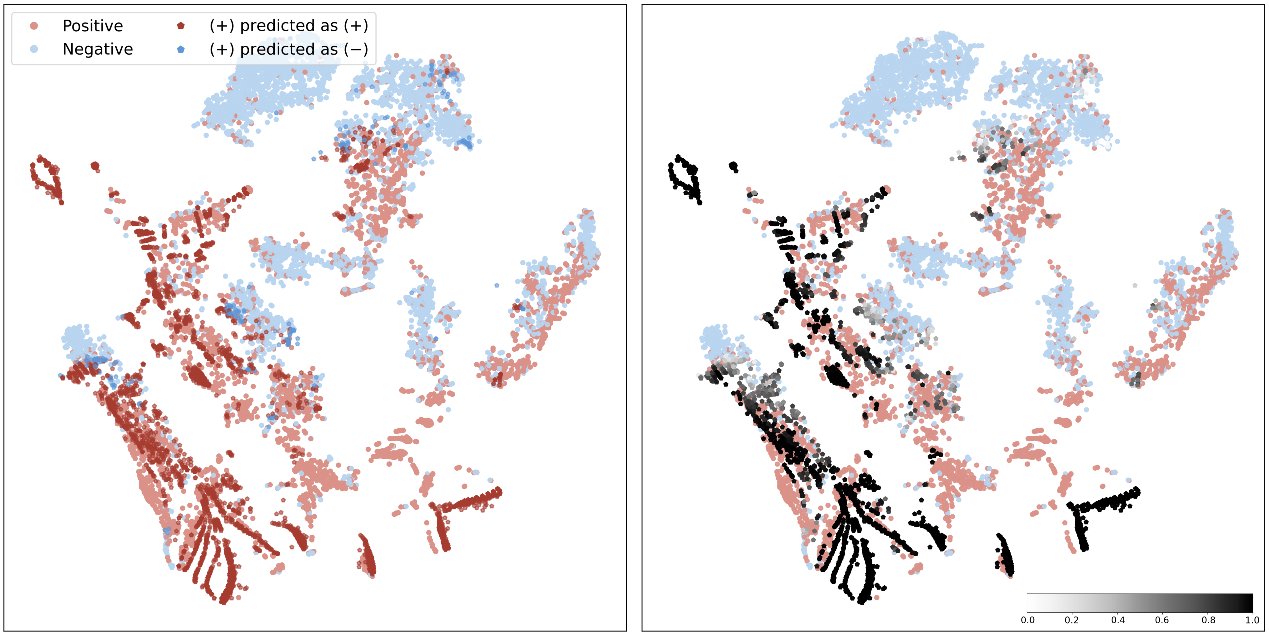


**Fig. 10.** t-SNE representation of the edge space. For both plots, Edges used during training labeled as positive (1) and negative (0) in lighter colors. **Left**. Evaluation edges in daker color using a threshold of 0.5. **Right**.Evaluation edges without applying threshold, using the ouput probability in gray scale. t-SNE parameters for both plots: perplexity = 150, niter= 1000, met = 'cosine', init = 'random'
